# Positive linkage between bacterial social traits reveals that homogeneous rather than specialized behavioral repertoires prevail in natural *Pseudomonas* communities

**DOI:** 10.1101/477471

**Authors:** Jos Kramer, Miguel Ángel López Carrasco, Rolf Kümmerli

## Abstract

Bacteria frequently cooperate by sharing secreted metabolites such as enzymes and siderophores. The expression of different ‘public good’ traits can be interdependent, and studies on laboratory systems have shown that such trait linkage affects eco-evolutionary dynamics within bacterial communities. Here, we examine whether linkage among social traits occurs in natural *Pseudomonas* communities by examining investment levels and correlations between five public goods: biosurfactants, biofilm components, proteases, pyoverdines, and toxic compounds. Our phenotypic assays involving 315 isolates from soil and freshwater communities revealed that their social trait expression profiles varied dramatically, and that correlations between traits were frequent, exclusively positive, and sometimes habitat-specific. Our results indicate that *Pseudomonas* communities are dominated by isolates lying on a continuum between a ‘social’ type producing multiple public goods, and an ‘asocial’ type showing low investment into social traits. This segregation into different social types could reflect local adaptation to different microhabitats, or emerge from competition between different (social) strategies. Moreover, our results show that isolates with specialized trait repertoires are rare, suggesting limited scope for the mutual exchange of different public goods between isolates. Overall, our work indicates that complex interdependencies among social traits influence the evolution of microbial lifestyles in nature.

## INTRODUCTION

Bacteria are social organisms. They form multicellular structures such as biofilms, communicate with each other via chemical signals, and engage in group-coordinated motility, resource acquisition and ‘chemical warfare’ against predators or competitors (West *et al.* 2007; Foster 2010). Many of these social behaviours involve the cooperative release of costly secondary metabolites. For instance, biofilm formation requires the production of structural polysaccharides, group-coordinated motility often involves the production of biosurfactants, and resource acquisition is based on the production of exo-enzymes and metal-chelating siderophores (Griffin, West and Buckling 2004; Nadell, Xavier and Foster 2009; Kearns 2010; Jousset *et al.* 2013). Once released, these compounds are – in principle – publicly accessible to cells in the vicinity. Such ‘public goods’ (*sensu* Kümmerli and Ross-Gillespie 2014) do therefore not only benefit (clonemates of) the producer, but can also affect other community members. The resulting interactions cover all aspects of bacterial life, ranging from ‘cheating’, where strains deficient for the production of a public good free-ride on the common pool of this resource (West *et al.* 2006; Özkaya *et al.* 2017) over inter-strain competition (Cordero *et al.* 2012; Bruce *et al.* 2017; Leinweber, Weigert and Kümmerli 2018) to division of labour between bacterial strains exchanging different types of public goods (van Gestel, Vlamakis and Kolter 2015; Dragoš *et al.* 2018). There is increasing evidence that such social interactions play a crucial role during host infections (Rumbaugh *et al.* 2009; Pollitt *et al.* 2014; Andersen *et al.* 2015; Granato *et al.* 2018), in various biotechnological and environmental processes (Bachmann *et al.* 2016; Hesse *et al.* 2018) and within the microbiome (Rakoff-Nahoum, Foster and Comstock 2016).

While many aspects of bacterial social interactions are well understood for individual traits, it is important to consider that the expression of different traits is often interdependent. Such dependencies can give rise to an additional level of complexity, as selection on one trait can have pleiotropic consequences for other traits (Jousset *et al.* 2009; Wilder, Diggle and Schuster 2011; Dandekar, Chugani and Greenberg 2012; Wang *et al.* 2015; Allen *et al.* 2016). For example, for *Pseudomonas* bacteria it was shown that a reduction of exo-protease production can increase sensitivity to predation (Jousset *et al.* 2009) and reduce metabolic abilities and toxin resistance (Dandekar, Chugani and Greenberg 2012; Wang *et al.* 2015), thereby compromising the spread of the non-producers. Especially when such linkage occurs between two or more public-good traits, the potentially resulting pleiotropic effects can have far-reaching consequences. For instance, positive linkage – where mutations reducing the production of one public good might also reduce the production of a second public good – could accelerate the loss of cooperation, and could hence favour the emergence of ‘supercheaters’ that simultaneously free-ride on multiple public goods (Granato *et al.* 2018). Conversely, negative linkage – where a reduced production of one public good might lead to an increase in the production of another public good – could promote the emergence of ‘mosaic producers’ with a limited public goods repertoire, which could potentially complement each other (Driscoll *et al.* 2011; Ross-Gillespie, Dumas and Kümmerli 2015).

While these studies on laboratory systems show that interdependent trait expression and the resulting constraints are key determinants of bacterial social interactions and strain co-existence, we know little on whether such interdependencies occur in natural bacterial communities and whether they have disruptive or stabilising effects on communities. Here, we tackle this question and investigate whether and how the production of five different public goods is correlated across 315 *Pseudomonas* isolates from eight soil and eight freshwater communities (18-20 isolates per community). In a first step, we screened each isolate for the production of: (i) pyoverdines, a highly variable class of siderophores used to scavenge iron (Meyer *et al.* 2008); (ii) proteases used to digest extracellular proteins (Diggle *et al.* 2007); (iii) biosurfactants used during group-motility and hydrocarbon uptake (Koch *et al.* 1991; Xavier, Kim and Foster 2011); (iv) structural components used to form biofilms (Nadell, Xavier and Foster 2009); and (v) toxic compounds used to fight predators and competitors (Jousset 2012). We then used our phenotypic data set to ask whether: (a) isolates vary in the extent to which they invest in the public goods; (b) there is a phylogenetic signal for trait expression, where more closely related isolates produce similar amounts of a particular public good; (c) production levels of different public goods correlate with one another, (d) isolates invest to similar extents in multiple traits (homogeneous producers) or show variation in their investments across traits (heterogeneous mosaic producers); and (e) trait association patterns differ between habitats.

## MATERIALS AND METHODS

### Sampling, isolation and identification of Pseudomonads

We sampled 16 *Pseudomonas* communities from soil and pond habitats located on the campus of the University of Zürich (47.40° N, 8.54° E), Switzerland. Specifically, we collected eight soil cores and eight water samples, and plated the bacteria contained therein on a medium selective for fluorescent pseudomonads. In each case, we then picked 20 *Pseudomonas* isolates at random (320 isolates in total), and verified their affiliation to the group of fluorescent pseudomonads by amplifying and sequencing part of the housekeeping gene *rpoD* (a commonly used phylogenetic marker for pseudomonads) (Mulet *et al.* 2009). The 315 isolates that could be unequivocally identified as pseudomonads were preserved as freezer stocks until their use in our experiments (see Butaitė *et al.* 2017) for details on the sampling, isolation and identification procedure).

### Phenotypic screening of natural isolates

We used standard phenotypic assays to screen all isolates for the production of the five public goods (see Supplementary Material for details). Before each assay, we grew our isolates from freezer stocks for 24h in rich medium, either in 50 ml Falcon tubes under shaking conditions (190 rpm) (for the protease assay), or in 96-well plates under static conditions (for all other assays). We quantified biosurfactant production as the extent of the biosurfactant-induced collapse of droplets of culture supernatant on a hydrophobic surface (Bodour and Miller-Maier 1998). Specifically, we grew isolates for 24h in a glucose-supplemented minimal medium, harvested the supernatant, spotted 5 µl supernatant-droplets on an oil-coated lid of a 96-well plate, and then measured the droplet surface area after one minute using the software ImageJ (Schneider, Rasband and Eliceiri 2012). We used the crystal violet assay to quantify biofilm formation as a proxy for the production of biofilm components (Savoia and Zucca 2007). In particular, we first grew isolates in Lysogeny Broth (LB) for 24h in 96-well plates that allow for the formation of surface-attached biofilms (round bottom plates; Sarstedt, Germany), and then measured the amount of crystal violet retained by the newly formed biofilm at 570 nm using an Infinite M200 Pro microplate reader (Tecan Trading AG, Switzerland). To quantify protease production, we spotted 1 µL aliquots of bacterial culture onto skim milk agar in 24-well plates, and then used ImageJ to measure the size of the proteolytic halo around colonies after 24h of growth (Jousset *et al.* 2013). To quantify pyoverdine production, we grew isolates in iron-limited casamino acids medium for 24h, and measured the natural fluorescence of pyoverdines in the culture supernatants (emission|excitation at 400|460 nm, respectively) (Kümmerli *et al.* 2009). Finally, for the production of toxic compounds we developed a supernatant feeding assay to assess to which extent isolates affect the growth of six reference strains (*P. aureofaciens* ATCC13985, *P. entomophila, P. putida* IsoF, *P.syringae* B728a, *P.protegens* CHA0, and *P. aeruginosa* PAO1). We first generated supernatant from each isolate after 24h growth in LB, then exposed the reference strains to each supernatant, and finally measured their growth in LB during 24h. We used the mean of the toxic effects on the six reference strains as measure of toxicity. For each assay, we grew isolates under static conditions at 28°C in fourfold replication, and measured the growth of the public-goods producing cultures (as final optical density at 600 nm, or – in the case of protease production – as colony size).

### Phylogenetic signal in public goods production

To explore the relationship between the phylogeny of our *Pseudomonas* isolates and their public goods production, we constructed maximum-likelihood phylogenetic trees based on partial rpoD gene sequences. We constructed separate trees for soil and pond habitats, based on, respectively, 149 and 148 (of the 158 and 157) isolates for which we obtained sequence lengths ≥ 600 bp. We used *P. aeruginosa* PAO1 as an outgroup, and included 20 characterised fluorescent pseudomonads as references to cover the broad phylogenetic diversity of our natural isolates (see (Butaite *et al.* 2017) for details on tree construction). Next, we mapped the production profiles of isolates onto the trees (using the web tool iTOL; Letunic and Bork 2016), and then calculated Blomberg’s K for each trait and habitat, to quantify the extent to which phylogeny predicts public good production profiles. K-values close to one reflect strong phylogenetic signals, suggesting that phylogenetic history is the main factor driving the trait covariance among isolates). Conversely, K-values close to zero reflect weak signals, suggesting that factors apart from phylogeny affect trait evolution (Blomberg, Garland and Ives 2003). Note that we ignored branch lengths for the calculation of the phylogenetic signal (cf. Butaitė *et al.* 2017, 2018), because the high variability of branch lengths observed among our isolates tends to lead to an underestimation of phylogenetic signals.

### Statistical analysis

All statistical analyses were conducted using R version 3.5.0 (www.r-project.org). We used a six-step procedure to investigate the production of – and correlation among – different public goods in our *Pseudomonas* communities. First, we scaled the measurements of public goods production relative to the corresponding measurements of the laboratory reference strain (*P. aeruginosa* PAO1), grown alongside the natural isolates. In the case of toxin production, values were scaled relative to the growth of the reference strains in the absence of our isolates’ supernatants. We then transformed the supernatant effect (by calculating [supernatant effect - 1]*[-1]) so that values above and below zero indicate growth inhibition (i.e. toxicity) and stimulation, respectively. In a second step, we checked for links between growth and public goods production to preclude that differences in growth lead to spurious correlations among social traits. As growth was positively correlated with the production levels of biofilm components (Spearman’s rank correlation: ρ = 0.364, p < 0.001), proteases (ρ = 0.254, p < 0.001) and pyoverdines (ρ = 0.710, p < 0.001), we calculated ‘per capita’ production levels of these traits by dividing each trait measurement by the corresponding growth measurement. Conversely, growth was not correlated with biosurfactant (ρ = - 0.030, p = 0.594) and toxic compound (ρ = 0.070, p = 0.215) production.

In the third step of our analyses, we transformed the measurements of all five social traits using a rank-based inverse normal (RIN) transformation (Bishara and Hittner 2012). This allowed us to investigate the production of and correlations among public good traits using parametric analyses. The transformation was necessary, as the untransformed data unequivocally yielded heavily-skewed residual distributions. In the fourth step, we used a nested-design permutational multivariate analysis of variance (PERMANOVA) to test whether the social profiles, including all five social traits, differed between soil and pond isolates. In the fifth step, we used linear mixed models to test for (habitat-specific) links between the social traits. We used the ‘false discovery rate’ (Benjamini and Hochberg 1995) to correct the p-values of correlations between pairs of social traits for multiple testing. Where appropriate, we included ‘community’ as a random effect to account for our nested sampling design. In the sixth and final step, we investigated to what extent isolates showed heterogeneous as opposed to homogeneous production profiles. For any given trait pair, we divided the area defined by the range of the observed traits values into two parts: a middle part enclosing the diagonal and comprising 50% of the total area, and two outer parts in the lower right and upper left corner, each comprising 25% of the total area (see Figure 3). Isolates falling outside or inside the middle part were classified as heterogeneous (score = 1) or homogeneous (score = 0) producers, respectively. For each isolate, we then averaged its scores for all trait pairs and tested whether the degree of heterogeneity observed among our isolates deviated from the null expectation. To obtain this null expectation, we simulated associations among five ‘random social traits’ with the same mean and standard deviation as the actual five social traits. Simulations involved 1000 independent runs, across which we averaged the simulated heterogeneity values. Further details on the statistical analyses and simulations are provided in the Supplementary Material.

## RESULTS

### Natural Pseudomonas isolates greatly vary in their social profiles

The social profile of our 315 *Pseudomonas* isolates did not differ between habitats (PERMANOVA: n_perm_ = 1000; habitat: F_1_ = 1.864, p = 0.108, Fig. 1a), but varied substantially among communities (F_14_ = 4.228, p = 0.001, Fig. 1b-f) due to community-level differences in the production of all five social traits (Table 1). Within-community variation followed trait-specific patterns (Table S1). In particular, communities were dominated by isolates producing no or low amounts of biosurfactants, while harbouring only few members producing high amounts of this public good (Fig. 1b). A similar albeit less pronounced pattern governed variation in biofilm formation, with most isolates producing low to intermediate amounts of biofilm components, and only few isolates investing more substantially in this social trait (Fig. 1c). Within-community variation in the production of proteases was characterized by a bimodal pattern, whereby 35.9 % of all isolates produced no proteases at all, while the remaining isolates produced considerable amounts of this public good (Fig. 1d). Variation in pyoverdine production spanned the entire continuum from zero to extremely high production levels (Fig. 1e). Similarly, the extent to which isolates had toxic effects on our references strains covered the entire continuum from being atoxic to strongly inhibitory (Fig. 1f). In summary, the high variation in the production of all five types of public goods leaves ample scope for the occurrence of both negative and positive trait associations.

**Table 1.**
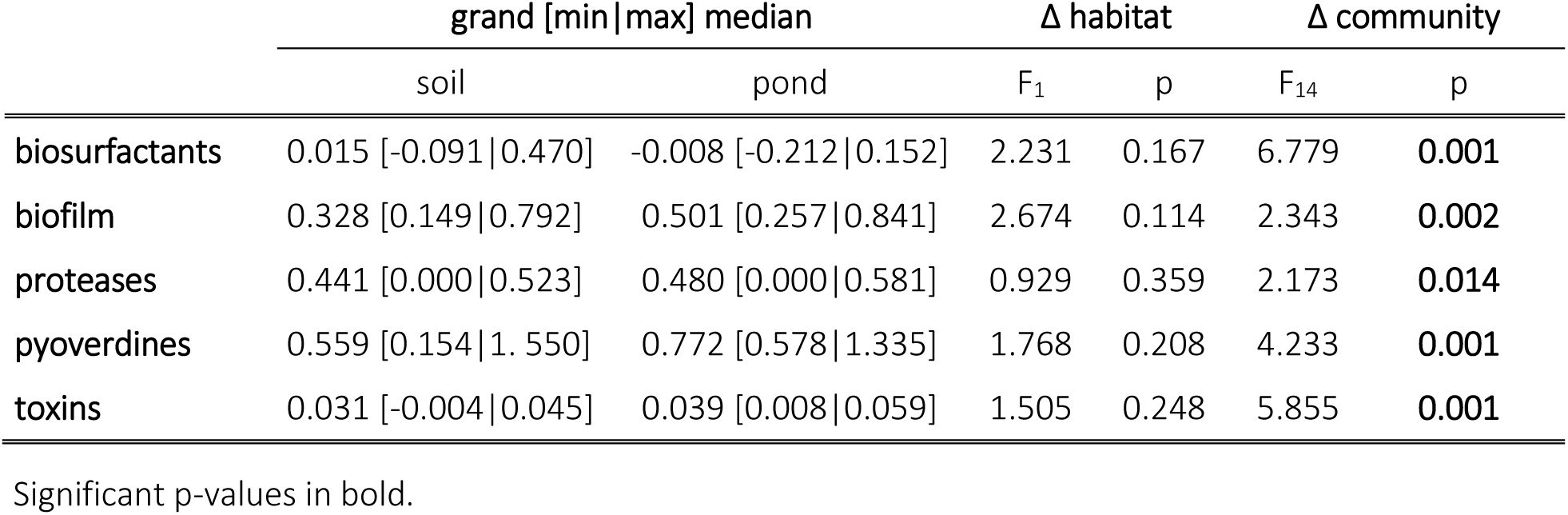
Among-community variation in the expression of public good traits. Given are the grand median as well as the minimal and maximal median of the production of biosurfactants, biofilm components, proteases, pyoverdines and toxic compounds in soil and freshwater communities, as well as the results of tests for habitat- and community-specific differences in the production levels of these traits.

| | grand [min\|max] median | | $\Delta$ habitat | | $\Delta$ community | |
| --- | --- | --- | --- | --- | --- | --- |
|  | soil | pond | F<sub>1</sub> | p | F<sub>14</sub> | p |
| **biosurfactants** | 0.015 [-0.091\|0.470] | -0.008 [-0.212\|0.152] | 2.231 | 0.167 | 6.779 | **0.001** |
| **biofilm** | 0.328 [0.149\|0.792] | 0.501 [0.257\|0.841] | 2.674 | 0.114 | 2.343 | **0.002** |
| **proteases** | 0.441 [0.000\|0.523] | 0.480 [0.000\|0.581] | 0.929 | 0.359 | 2.173 | **0.014** |
| **pyoverdines** | 0.559 [0.154\|1.550] | 0.772 [0.578\|1.335] | 1.768 | 0.208 | 4.233 | **0.001** |
| **toxins** | 0.031 [-0.004\|0.045] | 0.039 [0.008\|0.059] | 1.505 | 0.248 | 5.855 | **0.001** |

Significant p-values in bold.

**Figure 1.**
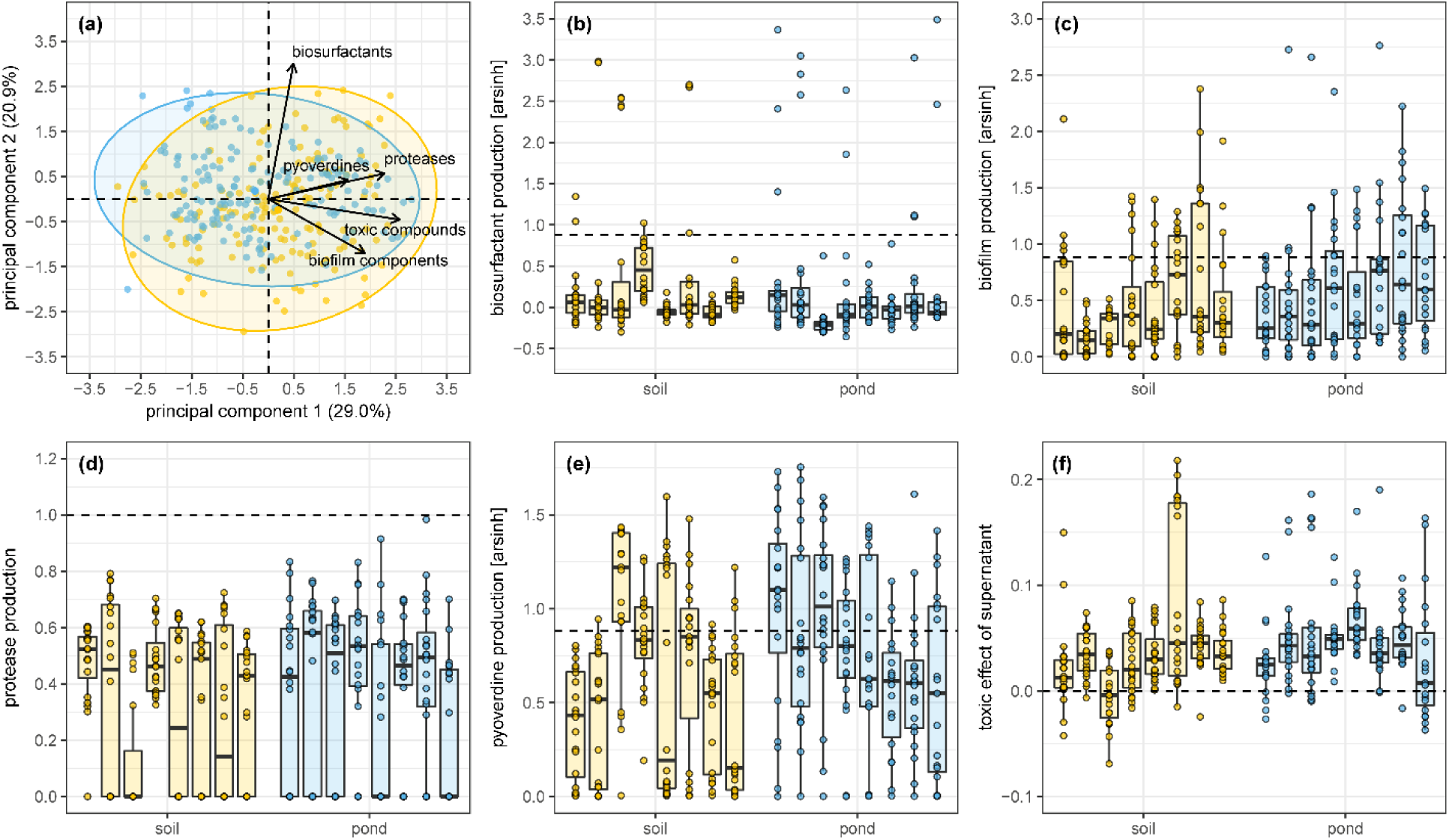
Social profile of *Pseudomonas* isolates. Depicted are (a) an overall representation of the social profile of 315 natural *Pseudomonas* isolates from eight soil and eight pond communities (in yellow and blue, respectively), as well as the variation in the production of (b) biosurfactants, (c) biofilm components, (d) proteases, (e) pyoverdines and (f) toxic compounds within and across communities. The depiction of the social profile in (a) is based on the first two components extracted from a principal component analysis based on the RIN-transformed values. Shaded areas are 95% confidence ellipses. The values of (b)-(e) are relative to the corresponding value of a laboratory control (*P. aeruginosa* PAO1, indicated by the dashed line). Values in (f) are the average toxic effect of the isolates’ supernatant on the growth of six reference strains (the dashed line indicates a neutral effect on growth).

### Phylogenetic signal for public goods production is higher in soil compared to pond

Our natural isolates covered an enormous phylogenetic diversity within the genus *Pseudomonas* (Fig. 2). This diversity affected social trait expression, as we detected a phylogenetic signal in the production levels of all five public goods (Table 2, Fig. 2a-b). Overall, phylogenetic signals were higher among soil than among freshwater isolates (paired t-test; t_4_ = 3.425, p = 0.027). In both habitats, the signals were significantly higher than expected from the null model for all traits, with the exception of biofilm formation among freshwater isolates (Table 2). Signals were generally of intermediate strength, indicating that both common ancestry and frequent changes in trait expression profiles within clades of related isolates shape public good production profiles.

**Table 2.** Phylogenetic signal in the production of public goods. Given are the phylogenetic signal (Blomberg’s K) and the results of a randomization test (under the null hypothesis of the absence of a phylogenetic signal) for the production of biosurfactants, biofilm matrix components, proteases, pyoverdines, and toxic compounds among soil and freshwater isolates, respectively.

|  | soil |  | freshwater |  |
| --- | --- | --- | --- | --- |
|  | Blomberg's K | p | Blomberg's K | p |
| biosurfactants | 0.378 | **0.001** | 0.203 | **0.024** |
| biofilm | 0.342 | **0.001** | 0.166 | 0.073 |
| proteases | 0.539 | **0.001** | 0.350 | **0.001** |
| pyoverdines | 0.390 | **0.001** | 0.382 | **0.001** |
| toxins | 0.283 | **0.001** | 0.211 | **0.005** |

Significant p-values in bold.

**Figure 2.**
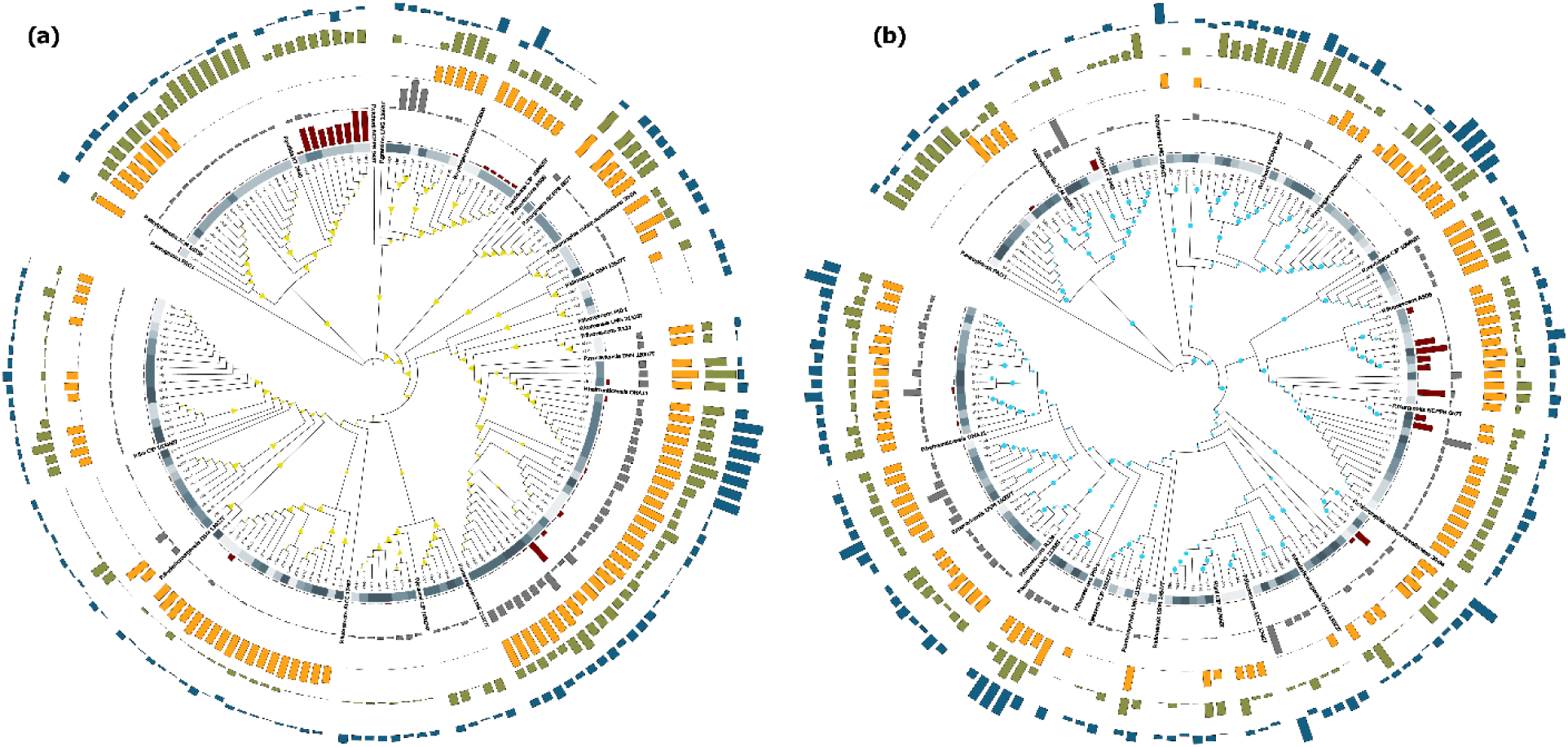
Maximum-likelihood cladograms for soil (a) and pond (b) isolates based on partial rpoD sequences. For both habitats, published *rpoD* sequences of 20 well-characterized fluorescent pseudomonads and *P. aeruginosa* PAO1 were integrated into the cladograms to demonstrate taxonomic affiliation and the high diversity of our environmental isolates. Yellow triangles and blue circles indicate bootstrap values (50-100%) for branches in the soil and the pond cladogram, respectively. From the inside outwards, bars depict the relative production levels of biosurfactants (red), biofilm components (grey), proteases (orange), pyoverdine (green), and toxic compounds (blue), respectively. Colour strips around cladograms represent the communities (in shades of grey) from which isolates originated.

### Positive correlations between public good traits dominate in both habitats

The production levels of the five examined public goods were positively correlated in seven out of ten possible pairwise comparisons (Table 3, Fig. 3). In particular, we observed habitat-independent positive correlations between the production of proteases and, respectively, biofilm components, pyoverdines, and toxic compounds, as well as between the production of pyoverdines and toxic compounds (Table 3, Fig. 3). There were also habitat-specific patterns: the production of proteases and biosurfactants correlated only among freshwater (slope ± se 0.181 ± 0.073, t_124.5_ = 2.474, p = 0.022) but not soil isolates (−0.053 ± 0.080, t_149.2_ = −0.654, p = 0.556), whereas the production of pyoverdines and biofilm components was positively correlated among soil (0.306 ± 0.070, t_153.4_ = 4.387, p < 0.001) but not freshwater isolates (−0.075 ± 0.083, t_154.7_ = −0.905, p = 0.432). Moreover, the production of biofilm components increased with toxicity in both habitats, but this increase was more pronounced among soil (0.385 ± 0.076, t_153.8_ = 5.104, p < 0.001) as compared to freshwater isolates (0.178 ± 0.072, t_152.3_ = 2.480, p = 0.022). Finally, we found that biosurfactant production was neither correlated with biofilm formation, nor with the production of pyoverdines and toxic compounds (Table 3, Fig. 3).

**Figure 3.**
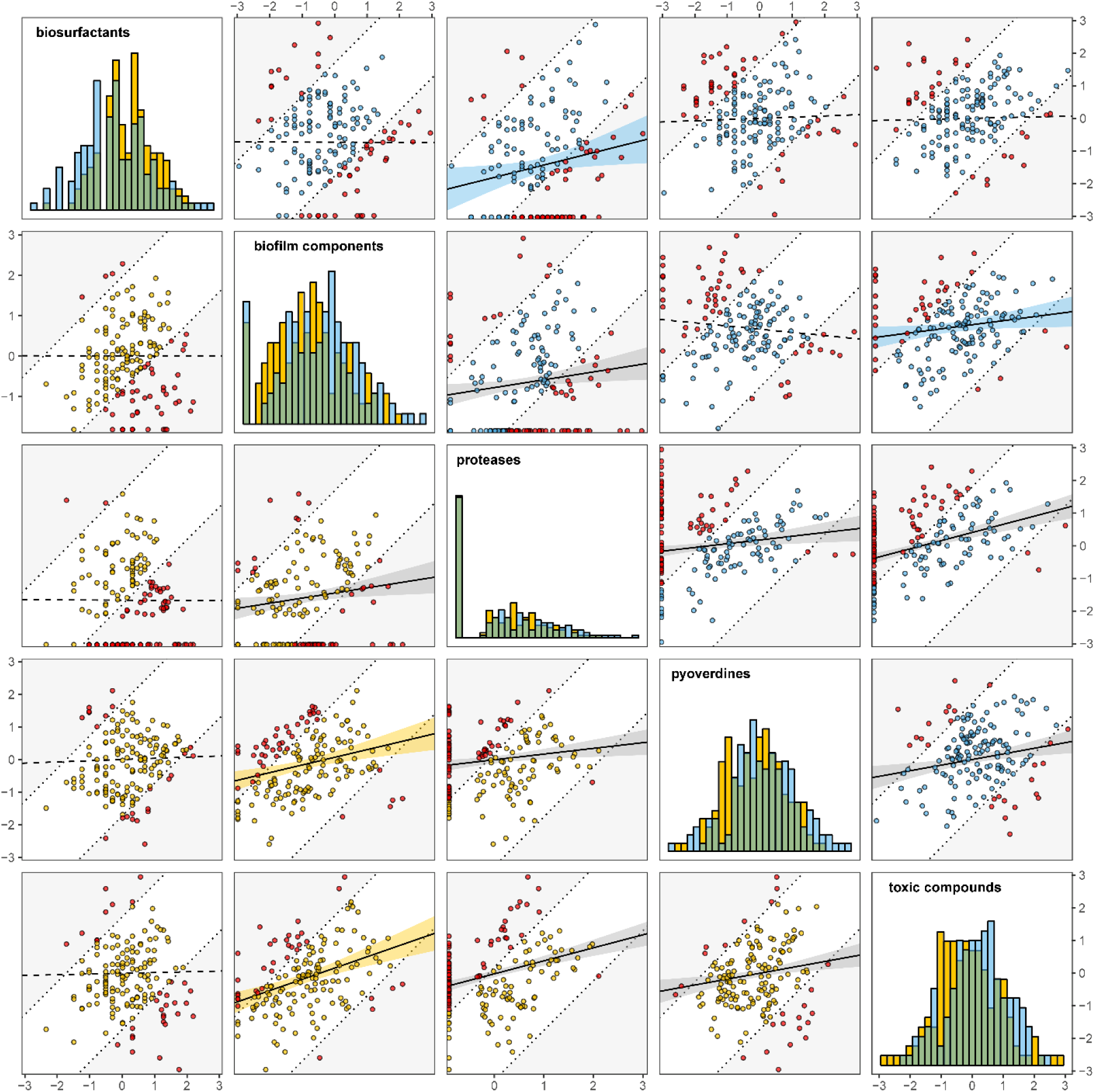
Correlation patterns between social traits. Shown are the pairwise correlations between social traits of 315 natural *Pseudomonas* isolates from eight soil (in yellow) and eight pond (in blue) communities. Solid and dashed lines depict significant and non-significant relationships, respectively. Bands in habitat-specific coloration as well as grey bands are 95 % confidence intervals derived from linear models and represent, respectively, habitat-specific correlations and correlations that are consistent across habitats. Isolates above the upper or below the lower dotted lines were classified as mosaic producers (in red).

**Table 3.** Pairwise correlations among public good traits. Shown are the (habitat-specific) pairwise correlations of the production of biofilm matrix components, proteases, pyoverdine and toxic compounds with the respective other social traits.

|  | biofilm |  | proteases |  | pyoverdines |  | toxins |  |
| --- | --- | --- | --- | --- | --- | --- | --- | --- |
| | $\chi^2$ | p | $\chi^2$ | p | $\chi^2$ | p | $\chi^2$ | p |
| **biosurfactants (bis)** | 0.140 | 0.708 | 7.438 | **0.006** | 2.108 | 0.191 | 2.109 | 0.191 |
| habitat | 2.833 | 0.092 | 1.195 | 0.274 | 1.976 | 0.160 | 1.746 | 0.186 |
| **bis : habitat** | 0.721<sup>§</sup> | 0.396<sup>§</sup> | 4.316 | **0.038** | 0.960<sup>§</sup> | 0.327<sup>§</sup> | 3.266<sup>§</sup> | 0.071<sup>§</sup> |
| **biofilm (bif)** |  |  | 6.690 | **0.021** | 0.577 | 0.447 | 6.372 | **0.012** |
| habitat |  |  | 0.487 | 0.485 | 1.324 | 0.250 | 0.921 | 0.337 |
| **bif : habitat** |  |  | 0.960<sup>§</sup> | 0.327<sup>§</sup> | 10.785 | **0.001** | 3.983 | **0.046** |
| **proteases (pro)** |  |  |  |  | 12.255 | **0.001** | 45.943 | **< 0.001** |
| habitat |  |  |  |  | 1.172 | 0.279 | 1.106 | 0.293 |
| **pro : habitat** |  |  |  |  | 3.689<sup>§</sup> | 0.055<sup>§</sup> | 0.112<sup>§</sup> | 0.738<sup>§</sup> |
| **pyoverdines (pvd)** |  |  |  |  |  |  | 21.689 | **< 0.001** |
| habitat |  |  |  |  |  |  | 0.735 | 0.391 |
| **pvd : habitat** |  |  |  |  |  |  | 3.109<sup>§</sup> | 0.078<sup>§</sup> |

<sup>§</sup> values prior to the removal of the interaction from the model. Significant p-values in bold. Note that we generally obtain the same results when the respective ‘response’ and the ‘explanatory’ social traits are swapped in the models testing for their correlation (swapping toxic compounds and biosurfactant production gives an additional positive correlation between these variables in the pond; see Table S2).

### Mosaic producers with heterogeneous social profiles are rare

The frequency of isolates with heterogeneous investment into pairs of social traits varied considerably, ranging from 15.9% (pyoverdines vs. toxic compounds) to 50.2% (proteases vs. biosurfactants), but was overall moderate, with heterogeneous producers representing 31.7 ± 12.1 % (mean ± SD) of all isolates. The average degree of heterogeneity across all five traits was lower than expected under the assumption of random trait expression (one-sample t-test; t_314_ = −2.192, p = 0.029, Fig. S1), indicating that mosaic producers with heterogeneous social profiles were rarer than isolates with homogeneous profiles.

## Discussion

Although linkage between bacterial social traits occurs frequently and has important consequences for strain interactions, pathogenicity and community functioning in laboratory systems (Harrison and Buckling 2009; Driscoll *et al.* 2011; Ross-Gillespie, Dumas and Kümmerli 2015; Granato *et al.* 2018), we know little about the prevalence and direction of social trait associations in natural bacterial communities. Here we addressed this open issue by examining and comparing the production levels of five social traits across 315 *Pseudomonas* isolates originating from eight soil and eight freshwater communities. The social traits we focused on included the production of biosurfactants used during group motility; matrix components required for biofilm formation; proteases used for extra-cellular protein digestion; pyoverdines required for iron scavenging; and toxic compounds used to fight competitors and predators. We found that (i) natural isolates varied dramatically in the extent to which they invest in these social traits, and showed that (ii) correlations between social traits occurred frequently, (iii) were exclusively positive, and (iv) in some cases habitat-specific. A comparison across traits suggest that natural *Pseudomonas* communities are dominated by isolates that lie on a continuum between two main phenotypes – a ‘social’ type that is proficient in expressing multiple social traits, and an ‘asocial’ type that generally shows low investment levels into any of the social traits (see also Ghoul *et al.* 2014). Conversely, mosaic producers with a specialized trait repertoire characterized by high investment in only some (but low investment in other) social traits were relatively rare. In the sections below, we offer potential mechanistic and evolutionary explanations for the observed patterns and discuss their implications.

At the mechanistic level, correlated trait expression could arise because public goods are linked at the regulatory level. Well-known and wide-spread co-regulatory systems in pseudomonads include quorum sensing (QS) systems, the GacS/GacA two-component system, and systems where public goods themselves serve as signals to co-regulate other social traits (Lamont *et al.* 2002; Jimenez *et al.* 2012; Dumas, Ross-Gillespie and Kümmerli 2013). Mechanisms of positive regulatory trait linkage such as QS could explain both the observed high investments into multiple public goods among ‘social’ types, and the low investment into public goods among ‘asocial’ types. The ‘asocial’ types could evolve through the acquisition of mutations in their QS-regulon, where the loss of one trait spurs the concomitant loss of other traits (Jousset *et al.* 2009; Dandekar, Chugani and Greenberg 2012; Wang *et al.* 2015). An alternative explanation is that the growth and/or the QS-threshold differs between isolates, meaning that some of the ‘asocial’ types might principally be able to express the social traits, but would only do so at cell densities higher than those achieved in our experiments. One way to explore whether this alternative explanation applies for our isolates is to test for non-linear relationships between trait investment and growth (as one would expect for QS-induced traits; Allen *et al.* 2016). We indeed observed such non-linear relationships for biofilm components, pyovedines, and toxic compounds, but not for the other traits (Fig. S2). To explore the consequences of these non-linear effects, we performed a second statistical analysis where we first fitted robust linear models with quadratic terms to test for trait-specific, non-linear relationships between production levels and growth, and then re-calculated all pairwise correlations between social traits using the rankit-transformed residuals extracted from these models (see the Supplementary Material for details). These analyses generally yielded results that match those shown in Fig. 3 (Table S3) with a few exceptions: we neither recovered the habitat-independent correlation between proteases and biofilm components, nor the pond-specific correlations between proteases and biosurfactants, and between pyoverdines and toxins. These findings suggest that certain correlations observed in Fig. 3 partly arise because of non-linear relationships between trait investment and growth, suggesting that QS-regulation or other density-dependent regulatory mechanism could be involved.

A further mechanistic explanation for correlated trait expression is environmental trait induction, which would suggest that isolates classified as non-producers for a particular social trait might actually express this trait under different environmental conditions. As discussed above, some isolates might grow poorly in our artificial media and thereby remain below the QS threshold required for social trait expression. Moreover, some isolates might only express certain social traits under specific nutrient conditions (Xavier, Kim and Foster 2011) or in response to relevant cues (Nadell, Xavier and Foster 2009; Dumas, Ross-Gillespie and Kümmerli 2013). Such environmental effects might explain the generally low levels of biosurfactant production, and thus the scarcity of correlations between the production of biosurfactants and the other public goods. However, it is unlikely that environmental trait induction explains all of our results, because it should lead to the seemingly random expression of traits across different media, and thus mask rather than yield correlations between traits.

At the evolutionary level, correlated trait expression could arise because high and low public good producers occupy – and are locally adapted to – different niches. In this scenario, the ‘asocial’ type would occur in microhabitats where social traits are not required or where coordinated actions among cells are difficult due to low bacterial densities (Kümmerli *et al.* 2014; Butaitė *et al.* 2018). Conversely, the ‘social’ type should mainly occur in microhabitats supporting higher bacterial densities and/or featuring conditions under which cooperation is beneficial. This would indicate that social traits are readily lost if not needed, and that conditions favouring the production of public goods typically lead to the evolution of generalists that invest homogeneously into the production of multiple public goods. Alternatively, ‘social’ and ‘asocial’ types might co-occur in the same microhabitats. In this case, that small differences in initial trait values could affect the complex feedback between multiple social traits, and thereby foster co-existence of different stable strategies at the population level (Brown and Taylor 2010). Similarly, spatio-temporal fluctuations in environmental conditions could alter the relative selective advantage of the two types and promote their co-existence (Or *et al.* 2007). Finally, social interactions involving the exploitation and cross-use of public goods among isolates could also explain the positive correlations between traits, whereby isolates investing nothing or little in public goods could act as ‘super-cheaters’ freeriding on the public goods produced by others (Diggle *et al.* 2007; Nadell, Xavier and Foster 2009; Van Gestel *et al.* 2014; Butaitė *et al.* 2017). At least for siderophores, cheater-cooperator interactions have been reported to occur in terrestrial and aquatic natural systems (Cordero *et al.* 2012; Bruce *et al.* 2017; Butaitė *et al.* 2017).

If social interactions drive the evolution of the ‘social’ and ‘asocial’ types, their co-existence can be explained in multiple ways. In the case of competitive interactions, where ‘asocial’ types exploit the ‘social’ types, public goods cooperation and isolate co-occurrence could be maintained by negative frequency-dependent selection (Ross-Gillespie *et al.* 2007). The concept of frequency-dependent selection predicts that cheaters only experience high relative fitness benefits when rare, but not when common, leading to an equilibrium frequency at which the two isolate types can co-exist. Antagonistic co-evolution between the two types could also explain the co-existence of the asocial and social types (Kümmerli *et al.* 2015; O’Brien *et al.* 2017). For instance, increased biofilm formation in isolates that invest highly in proteases and pyoverdine could be a direct response to cheating, because biofilm formation induces spatial segregation between cell types (Van Gestel *et al.* 2014; Irie *et al.* 2017) and can thus exclude cheaters from the efficient exploitation of these public goods (Drescher *et al.* 2014). In the case of commensalistic interactions, isolate co-existence could be favoured if a certain amount of public goods is required to support the community as predicted by the Black Queen Hypothesis (Morris, Lenski and Zinser 2012). This hypothesis posits that the utilization of public goods by the ‘asocial’ type does not affect the fitness of the ‘social’ type (Morris, Lenski and Zinser 2012) – an assumption that might not be met in our communities, where isolates are generally closely related and therefore presumably have overlapping resource requirements. In the case of mutualistic interactions, isolate co-existence could arise through specialization, where the different public goods are exchanged between isolates (Dragoš *et al.* 2018). However, the relative rarity of mosaic producers with specialised trait repertoires suggest that cross-feeding scenarios, like those described in laboratory systems (Driscoll *et al.* 2011; Pande *et al.* 2014; Kim, Levy and Foster 2016; Dragoš *et al.* 2018), might be relatively rare among our natural isolates.

Our observation that there are habitat-specific correlations suggest that trait linkage and the underlying mechanisms can change in response to different environmental selection pressures. Changes in the local environment, for instance when a soil isolate is washed into a pond, could thus rapidly promote the breaking of existing but superfluous linkage patterns or select for new ones that are beneficial in the new habitat (Dos Santos, Ghoul and West 2018). The moderate phylogenetic signals we detected are in line with this notion, as they suggest that the expression levels of social traits can evolve relatively flexibly. In this context, the overall higher phylogenetic signal in soil as compared to freshwater communities could indicate faster and more flexible evolution of public good repertoires in ponds.

In conclusion, our findings indicate that social trait linkage (i) is common in natural *Pseudomonas* communities, (ii) has more likely evolved as a result of competitive rather than commensalistic or mutualistic interactions (Foster and Bell 2012; Oliveira, Niehus and Foster 2014), and (iii) underlies the frequent occurrence of isolates with homogeneous behavioral repertoires lying on a continuum between two main isolate types – a ‘social’ type that is proficient in expressing multiple social traits, and an ‘asocial’ type that generally shows low investment levels into most of the social traits. While our study solely focuses on pseudomonads, it would be interesting to see whether the same patterns of trait linkage occur beyond the genus boundary, especially for traits such as proteases and toxins, which have more broad-spectrum effects than pyoverdine, which is a characteristic trait of the fluorescent pseudomonads. Moreover, while our use of standardized laboratory assays in a ‘common garden’ laboratory environment allowed us to assess the isolates’ genetic potential to produce specific public goods, we do not know to which extent they would do so under natural conditions, and whether isolates can plastically adjust public goods production in response to changes in environmental or social conditions. We also know little about the dynamics and change of isolate frequency over time since we sampled only a single snapshot in time. These points highlight that our work only provides a first glimpse of the complexity of microbial life in natural communities. This glimpse, however, suggests that social trait linkage could play an important role in explaining community composition and functioning.

## Authors’ contributions

JK and RK designed the experiments; JK and MALC conducted the experiments; JK performed the statistical analyses; JK, MALC and RK wrote the paper. All authors gave final approval for publication.

## Acknowledgements

We thank Elena Butaitė for collecting the natural isolates, and Stefan Wyder for help with the phylogenetic analysis.

