## Supplementary Material for "Positive linkage between bacterial social traits reveals that homogeneous rather than specialized behavioral repertoires prevail in natural *Pseudomonas* communities"

This file contains the following Supplementary Information:

- **Supplementary Methods**
- **Supplementary Analyses**
- **Supplementary References**
- **Supplementary Tables S1-S3**
- **Supplementary Figure S1+S2**

### Supplementary Methods

Unless otherwise stated, all cultures were incubated at 28°C under static conditions. With the exception of the biofilm assay (see below), we used non-tissue-culture-treated 96-well plates for all measurements of public good traits. These plates hamper bacterial surface attachment. Each of the plates featured multiple blanks and contained all isolates of one community and our laboratory reference strain *P. aeruginosa* PAO1 in fourfold replication. All liquid cultures were extensively mixed before photometric measurements, and we always obtained multiple photometric measurements per well.

**Measurement of biosurfactant production.** We quantified biosurfactant production using a 'modified drop-collapse assay' that involves the measurement of the biosurfactant-dependent spread of droplets of bacterial supernatant on a hydrophobic surface. Specifically, we first grew single isolates from freezer stocks in 200 µl Lysogeny Broth (LB) for 24 h, and then diluted the resulting cultures 100-fold in 200 µl mineral salts medium (details in (Bodour and Miller-Maier 1998)). After another incubation step of 24 h, we measured the growth (optical density, measured at 600 nm [$OD_{600}$]) of each culture using an Infinite M200 Pro microplate reader (Tecan Trading AG, Switzerland), and then quantified the amount of biosurfactants in the supernatant. To this end, we centrifugated the cultures at 3600 rcf for 10 min (Type 5804R, Eppendorf, Germany), and then delivered 5 µl aliquots of the supernatant on the lid of a 96-well plate that had been coated with light mineral oil (M8410, Sigma-Aldrich, USA). On this hydrophobic surface, droplets of supernatant spread out [or bead up] to a size that is proportional to the amount of biosurfactants [or highly-polar and soluble compounds] in the supernatant (Bodour and Miller-Maier 1998). To obtain a measure of biosurfactant production, we took photographs of the supernatant-droplets one minute after their application using a standard digital camera, and then measured their surface area using the software *ImageJ* (Schindelin *et al.* 2012; Schneider, Rasband and Eliceiri 2012).

**Measurement of biofilm formation.** We quantified the production of biofilm components by measuring the amount of crystal violet retained by surface-attached biofilms (Savoia and Zucca 2007). In particular, we first grew single isolates from freezer stocks for 24 h in 200 µl LB. We then diluted each of the resulting cultures 50-fold in 100 µl LB and incubated them for another 24 h in 96-well cell culture plates that allow for the formation of surface-attached biofilms (round bottom plates; Sarstedt, Germany). Subsequently, we transferred the growth medium of each plate to a fresh plate, and measured the growth ($OD_{600}$) of the planktonic cells contained therein. In the meantime, we added 100 µl crystal violet (0.1%) to each well of the original plate (which potentially contained surface-attached biofilm cells) and incubated it for 30 min to allow for biofilm staining. Subsequently, we washed the plate thoroughly with $ddH_20$, dried it at room temperature for 30 min, and then added 120 µl of Dimethyl sulfoxide (DMSO) to each well to solubilize the biofilm-bound crystal violet in a final incubation step of 20 min. Finally, we measured the production of biofilm matrix components as the $OD_{570}$ of the crystal violet solution.

**Measurement of protease production.** We quantified the production of (exo-) proteases by measuring the size of the proteolytic halo formed around protease-producing colonies growing on skim milk agar plates (Jousset *et al.* 2013). Specifically, we grew single isolates from freezer stocks at 28°C and 190 rpm in 50 ml Falcon tubes containing 5 ml LB, respectively. After 24 h, we washed the resulting cultures twice, and then diluted them to a standardized optical density ($OD_{600} = 1$) in 0.85% NaCl. Subsequently, we transferred aliquots of 1 µl of each standardized culture into single wells of 24-well plates containing solidified skim milk agar (4% [m/V] skim milk powder, 5 g/L LB, and 15 g/L agar). We prepared six plates per community that each contained all isolates of a given community, the laboratory reference strain (*P. aeruginosa* PAO1), and multiple blanks. Following 24h of incubation, we took photographs of the plates with a standard digital camera, and then used *ImageJ* to measure bacterial growth (surface of the colony) and the production of secreted proteases (surface of the halo including the colony).

**Measurement of pyoverdine production.** We measured pyoverdine production by quantifying the natural fluorescence of this siderophore class in liquid culture (Kümmerli *et al.* 2009). Specifically, we first grew single isolates from freezer stocks for 24 h in 200 µl of an 'overnight medium' (Özkaya *et al.* 2018). Subsequently, we diluted each of these cultures 100-fold in 200 µl iron-limited casamino acids medium (containing 5 g casamino acids, 1.18 g $K_2HPO_4 \cdot 3H_2O$, and 0.25 g $MgSO_4 \cdot 7H_2O$ per liter) that had been supplemented with 25 mM HEPES buffer, 20 mM $NaHCO_3$ and 100 µg/ml human apo-transferrin (a strong natural iron chelator). After 24 h of incubation, we determined growth ($OD_{600}$) and then extracted 100 µl of supernatant by centrifugating the cultures at 2250 rcf for 15 min. Finally, we used this supernatant to measure the production of pyoverdines based on their natural fluorescence (relative fluorescence units [RFU] with excitation and emission at 400 nm and 460 nm, respectively).

**Measurement of toxin production.** We quantified the production of growth-inhibiting compounds by comparing the growth of five natural *Pseudomonas* strains (*P. aureofaciens* ATCC13985, *P. entomophila*, *P. putida* IsoF, *P.syringae* B728a, and *P.protegens* CHA0) and our laboratory control (*P. aeruginosa* PAO1) in the presence and absence of supernatant of our natural isolates. To generate this supernatant, we grew each isolate from its freezer stock in 200 µl LB for 24 h, respectively. Subsequently, we diluted each of the resulting cultures 40-fold into 200 µl LB. After 24 h of incubation, we measured the growth ($OD_{600}$) of the resulting cultures, centrifugated them at 2250 rcf for 10 min, and then purified the resulting supernatant by spinning it through a 3 µm glass fiber/0.2 µm Supor membrane at 1200 rcf (using an AcroPrep Advance 96-well filter plate [Pall Corporation, USA] attached to a 2.2 ml deep 96-well plate). The resulting cell-free supernatant was sealed in collection plates (using SILVERseal sealing mats; Greiner Bio-One, Austria) and stored at -20°C until further use. To test the effect of the supernatants on the growth of the six reference strains, we first grew these strains from freezer stocks for 24 h in 24-well plates containing 1.6 ml LB per well. We then inoculated each reference strain at a final $OD_{600} = 0.005$ in 96-well plates containing 180 µl LB and 20 µl of either supernatant or 0.85% NaCl per well. The NaCl had been subject to the same treatment as the supernatants (i.e. filtering and freezing), and either served as a blank or a negative control (inoculated with the reference strain) mimicking the addition of 'used-up' medium. After a last incubation step of 24 h, we measured the growth ($OD_{600}$) of the reference strains.

**Statistical analysis.** We used a six-step procedure to investigate the production of – and correlation among – different public goods in our natural *Pseudomonas* communities. First, we scaled the (blank-corrected) measurements of public goods production relative to the corresponding measurements of the laboratory reference strain that had been grown alongside the natural isolates, and then computed the median across replicates. To obtain a measure of an isolates average toxicity, we first scaled each reference strain's growth in the presence of a specific supernatant relative to its growth in the absence of that supernatant, and then calculated the arithmetic mean of this supernatant effect across the six reference strains. Finally, we transformed the supernatant effect (by calculating [X-1]*[-1], where X is the supernatant effect) so that values above and below zero indicate, respectively, an inhibition of growth (i.e. toxicity) and stimulation of growth. To preclude that differences in growth lead to spurious correlations among social traits, we checked for associations between growth (measured as final $OD_{600}$ of the public goods-producing culture, or – in the case of protease production – as colony size) and public goods production in a second step. As growth was positively correlated with the production levels of biofilm components (Spearman's rank correlation: $\rho = 0.364$, $p < 0.001$), proteases ($\rho = 0.254$, $p < 0.001$) and pyoverdines ($\rho = 0.710$, $p < 0.001$), we calculated 'per capita' production levels of these traits by dividing each trait measurement by the corresponding measurement of growth. Growth was independent of the production levels of biosurfactants ($\rho = -0.030$, $p = 0.594$) and toxic compounds ($\rho = 0.070$, $p = 0.215$). In the third step, we transformed the measurements of all five social traits into 'rankits', as preliminary analyses based on the untransformed measurements yielded heavily-skewed residual distributions. Rankits result from a rank-based inverse normal (RIN) transformation that involves converting the data into ranks, then converting these ranks into probabilities, and finally converting these probabilities into an approximately normal shape using the inverse cumulative normal function (Bliss 1967; Bishara and Hittner 2012). This transformation allowed us to investigate the production of and correlations among public good traits using parametric analyses.

In the fourth step of our statistical procedure, we tested whether the social profile (i.e. the expression of all five social traits) differed between soil and pond isolates. To this end, we performed a nested-design permutational multivariate analysis of variance (PERMANOVA) in which we used the five social traits as continuous response variables and the habitat as explanatory variable. Additionally, we included 'community-ID' as a co-factor to account for potentially confounding differences among communities within the same habitat. In the fifth step, we used linear mixed models to test for (habitat-specific) links between the social traits. Each of these models included one social trait as (continuous) response, as well as a second social trait (continuous), the habitat (soil or pond; categorical), and their interaction as explanatory variables. To account for our nested sampling design, we included 'community' as a random effect in each model. We corrected the p-values of the correlations between pairs of social traits for multiple testing using the 'false discovery rate' (Benjamini and Hochberg 1995). All statistical analyses were conducted using the statistics software R version 3.5.0 (www.r-project.org). Mixed models were implemented in R using the 'lmer'-function of the *lme4* package. The p-values of effects in these models were obtained using the 'Anova'-function of the *car* package. Blomberg's K was calculated using the 'phyloSignal'-function of the *phyloSignal* package. We used the 'nested.anova.dbrda'-function of the *BiodiversityR* package and the 'PCA'-function of the *FactoMineR* package to perform, respectively, the PERMANOVA and the principal component analysis used to depict the social profile.

In the sixth and final step, we investigated to what extent isolates showed heterogeneous rather than homogeneous production profiles. To this end, we scored for each pairwise combination of social traits whether an isolate showed a heterogeneous trait investment (yes = 1, no = 0), then averaged across the scores of all trait pairs, and finally tested whether the degree of heterogeneity observed among our isolates deviated from the null expectation (Figure S1). To be able to score an isolate's heterogeneity of investment into a given trait pair, we divided the area defined by the range of the two traits into two parts: a middle part enclosing the diagonal and comprising 50% of the total area, and two outer parts in the lower right and upper left corner, each comprising 25% of the total area. Isolates falling outside of [into] the middle part showed a dissimilar [similar] investment into the two traits, and were scored as heterogeneous [homogeneous] producers. We obtained the expected degree of heterogeneity in three steps. First, we created five 'random social traits', each with the same mean and standard deviation as one of the five actual social traits. In the next step, we assessed the number of randomly occurring heterogeneous producers for all possible pairs of the random variable, thereby mirroring our procedure for the determination of the actual number of heterogeneous producers in the pairs of actual social traits. Finally, we calculated the degree of heterogeneity across all pairs of random variables. We obtained the expected degree of heterogeneity by repeating the above procedure 1000 times.

### Supplementary Analyses

In the analyses presented in the main text, we tested for correlations between different social traits using 'per capita' (e.g. OD-corrected) values of trait production. Social traits in bacteria can be co-regulated via quorum sensing, and such regulation often results in a non-linear relationship between trait investment and growth. We ran an additional set of analyses to explore these effects and their impact on the results and conclusions presented in the main text. In a nutshell, we first fitted robust linear models with quadratic terms to test for trait-specific links between production levels and growth, and then re-calculated all pairwise correlations between social traits using the growth-corrected (rankit-transformed) residuals extracted from these models. We provide details on this alternative statistical procedure below, and outline its results and drawbacks.

**Statistical procedure.** To test for trait-specific relationships between social trait investment and growth, we fitted three robust linear models (via the 'lmrob' function of the *robustbase* package in R) on all five social traits. In each of these models, we entered the production values of one social trait as response variable, and the corresponding measurements of growth (with both a linear and a quadratic term) as explanatory variable. Robust models discount the influence of extreme values on parameter estimates (Maronna *et al.* 2019), and we used them here because our measurements of social traits featured many such values (see Fig. 1 in the main text). In the next step, we extracted the residuals from those models that revealed a link between trait values and growth (see below), thus obtaining trait values that were fully independent of growth (biosurfactant production: $\rho = -0.030$, $p = 0.594$; biofilm formation: $\rho = -0.087$, $p = 0.124$; protease production: $\rho = 0.023$, $p = 0.679$; pyoverdine production: $\rho = 0.007$, $p = 0.904$; toxin production: $\rho = 0.033$, $p = 0.562$). Finally, we transformed these values to rankits, and re-calculated the pairwise correlations between traits as well as the degree of heterogeneity following the procedure outlined in the main text and in the Supplementary Methods above.

**Results.** We found evidence for a non-linear increase of trait investment with growth in the case of biofilm formation (slope ± se; linear term: 1.498 ± 0.256, $t = 5.853$, $p < 0.001$; quadratic term: -0.553 ± 0.254, $t = -2.179$, $p = 0.030$; Fig. S2b), the production of pyoverdines (linear term: 4.626 ± 0.305, $t = 15.174$, $p < 0.001$; quadratic term: -2.181 ± 0.303, $t = -7.203$, $p < 0.001$; Fig. S2d), and the production of toxic compounds (linear term: 0.004 ± 0.031, $t = 0.137$, $p = 0.891$; quadratic term: -0.122 ± 0.031, $t = -3.997$, $p < 0.001$; Fig. S2e). In all three cases, trait production initially increased with growth, but this increase was attenuated (and ultimately reversed at least in the case of pyoverdine and toxic compounds) for higher growth values (Fig. S2). By contrast, the increase of protease production with growth was uniformly linear (linear term: 0.212 ± 0.060, $t = 3.555$, $p < 0.001$; quadratic term: -0.166 ± 0.299, $t = -0.556$, $p = 0.578$; Fig. S2c), whereas the production of biosurfactants was independent of growth (linear term: -0.018 ± 0.023, $t = -0.782$, $p = 0.435$; quadratic term: 0.024 ± 0.159, $t = 0.151$, $p = 0.880$; Fig. S2a). When using the residuals of these models (or, in the case of biosurfactants, the 'raw' trait values) in calculating the pairwise correlations between the five social traits, we obtained results similar to those presented in the main text. In particular, we found that the production levels of these traits were positively correlated in five out of ten possible pairwise comparisons (Table S3). We observed habitat-independent positive correlations between the production of proteases and, respectively, pyoverdines and toxic compounds, as well as between the production of biofilm components and toxic compounds (Table S3). In two additional cases, we found habitat-specific patterns (Table S3). Among soil isolates, the production of pyoverdines increased with biofilm formation (0.186 ± 0.063, $t_{154.4} = 2.977$, $p = 0.007$) and the production of toxic compounds (0.382 ± 0.093, $t_{155.9} = 4.116$, $p < 0.001$), whereas no such link was detectable among pond isolates (biofilm components: -0.091 ± 0.091, $t_{154.3} = -0.992$, $p = 0.428$; toxic compounds: 0.048 ± 0.067, $t_{154.1} = 0.726$, $p = 0.511$). Finally, we found no correlation between the production of biofilm components and proteases. Similarly, the production of biosurfactants was not correlated to any of the other social traits (Table S3).

Overall, the degree of heterogeneity in the expression profiles of our natural isolates was lower than expected under the assumption of random trait expression ($t_{314}$ = -2.088, p = 0.038).

**Table S1 | Within-community variation in the expression of public good traits.** Given are the median interquartile range (IQR; a measure of statistical dispersion) as well as the minimal and the maximal IQR of the production of biosurfactants, biofilm components, proteases, pyoverdines and toxic compounds within soil and freshwater *Pseudomonas* communities.

| | **median [min\|max] IQR** | |
|---|---|---|
| | soil | freshwater |
| biosurfactants | 0.201 [0.061\|0.591] | 0.201 [0.099\|0.364] |
| biofilm | 0.567 [0.203\|1.597] | 0.725 [0.477\|1.317] |
| proteases | 0.526 [0.147\|0.681] | 0.496 [0.146\|0.659] |
| pyoverdines | 0.764 [0.383\|1.540] | 0.867 [0.420\|1.177] |
| toxic compounds | 0.034 [0.022\|0.163] | 0.028 [0.013\|0.069] |

**Table S2 | Reversed pairwise comparisons among public good traits**. Shown are the (habitat-specific) pairwise correlations of the production of (A) biosurfactants, (B) biofilm components, (C) proteases, and (D) pyoverdines with the other social traits. The ‘response’ and the ‘explanatory’ traits are swapped to showcase the equivalency of the resulting correlations with those presented in Table 3 of the main text. [§]values prior to the removal of the interaction from the model. Significant p-values in bold. [$]the interaction reflects an increase of biosurfactant production with toxin production among pond (slope ± se: 0.206 ± 0.087, $t_{153.9}$ = 2.354, p = 0.030), but not among soil (-0.034 ± 0.067, $t_{156.0}$ = -0.508, p = 0.662) isolates. The corresponding interaction presented in the main text is marginally non-significant.

| | (A) biosurfactants | | (B) biofilm | | (C) proteases | | (D) pyoverdines | |
|---|---|---|---|---|---|---|---|---|
| | $\chi_1^2$ | p | $\chi_1^2$ | p | $\chi_1^2$ | p | $\chi_1^2$ | p |
| **toxins (tox)** | 6.769 | **0.009** | 27.486 | **< 0.001** | 45.288 | **< 0.001** | 20.783 | **< 0.001** |
| habitat | 2.432 | 0.119 | 1.306 | 0.253 | 0.084 | 0.773 | 0.655 | 0.418 |
| **tox : habitat** | 4.780 | **0.029**[$] | 2.198[§] | 0.138[§] | 1.496[§] | 0.221[§] | 0.365[§] | 0.546[§] |
| **pyoverdines (pvd)** | 2.217 | 0.197 | 0.832 | 0.362 | 10.944 | **0.002** | | |
| habitat | 2.490 | 0.115 | 1.979 | 0.160 | 0.343 | 0.558 | | |
| **pvd : habitat** | 0.693[§] | 0.405[§] | 12.903 | **< 0.001** | 2.767[§] | 0.096[§] | | |
| **proteases (pro)** | 8.508 | **0.004** | 6.762 | **0.017** | | | | |
| habitat | 2.478 | 0.115 | 2.169 | 0.141 | | | | |
| **pro : habitat** | 5.840 | **0.016** | 1.718[§] | 0.190[§] | | | | |
| **biofilm (bif)** | 0.113 | 0.737 | | | | | | |
| habitat | 2.308 | 0.129 | | | | | | |
| **bif : habitat** | 0.833[§] | 0.361[§] | | | | | | |

**Table S3 | Pairwise comparisons among public good traits based on (growth-corrected) residuals.** Shown are the (habitat-specific) pairwise correlations of the production of biofilm matrix components, proteases, pyoverdine and toxic compounds with the respective other social traits. All trait values are residuals extracted from robust linear models testing for a link between the production levels of a particular social trait and growth (see the Supplementary Analyses section for details). § values prior to the removal of the interaction from the model. Significant p-values in bold.

| | biofilm | | proteases | | pyoverdines | | toxins | |
|---|---|---|---|---|---|---|---|---|
| | $\chi_1^2$ | p | $\chi_1^2$ | p | $\chi_1^2$ | p | $\chi_1^2$ | p |
| **biosurfactants (bis)** | 0.614 | 0.473 | 2.109 | 0.251 | 0.984 | 0.428 | 1.496 | 0.379 |
| habitat | 2.129 | 0.145 | 3.021 | 0.082 | 0.680 | 0.409 | 2.336 | 0.126 |
| **bis : habitat** | 0.485§ | 0.486§ | 2.835§ | 0.092§ | 1.389§ | 0.239§ | 3.578§ | 0.059§ |
| **biofilm (bif)** | | | 0.147 | 0.701 | 0.965 | 0.326 | 20.016 | **< 0.001** |
| habitat | | | 2.373 | 0.124 | 0.467 | 0.495 | 1.518 | 0.218 |
| **bif : habitat** | | | 3.280§ | 0.070§ | 5.570 | **0.018** | 0.575§ | 0.448§ |
| **proteases (pro)** | | | | | 19.197 | **< 0.001** | 12.259 | **0.001** |
| habitat | | | | | 0.193 | 0.660 | 1.478 | 0.224 |
| **pro : habitat** | | | | | 0.546§ | 0.460§ | 0.720§ | 0.396§ |
| **pyoverdines (pvd)** | | | | | | | 0.605 | 0.437 |
| habitat | | | | | | | 1.365 | 0.243 |
| **pvd : habitat** | | | | | | | 7.852 | **0.005** |

**Figure S1 | The heterogeneity in social trait expression among natural *Pseudomonas* isolates.** Isolates with a low degree of heterogeneity invest to a similar extent in multiple social traits, whereas isolates with a high degree of heterogeneity show striking variation in their investment across traits (i.e. they have, respectively, homogeneous or heterogeneous production profiles). We found that the degree of heterogeneity (grey bars) varied widely across our 315 natural isolates, but was on average (red line) lower than expected under the assumption of random trait expression (dashed black line).

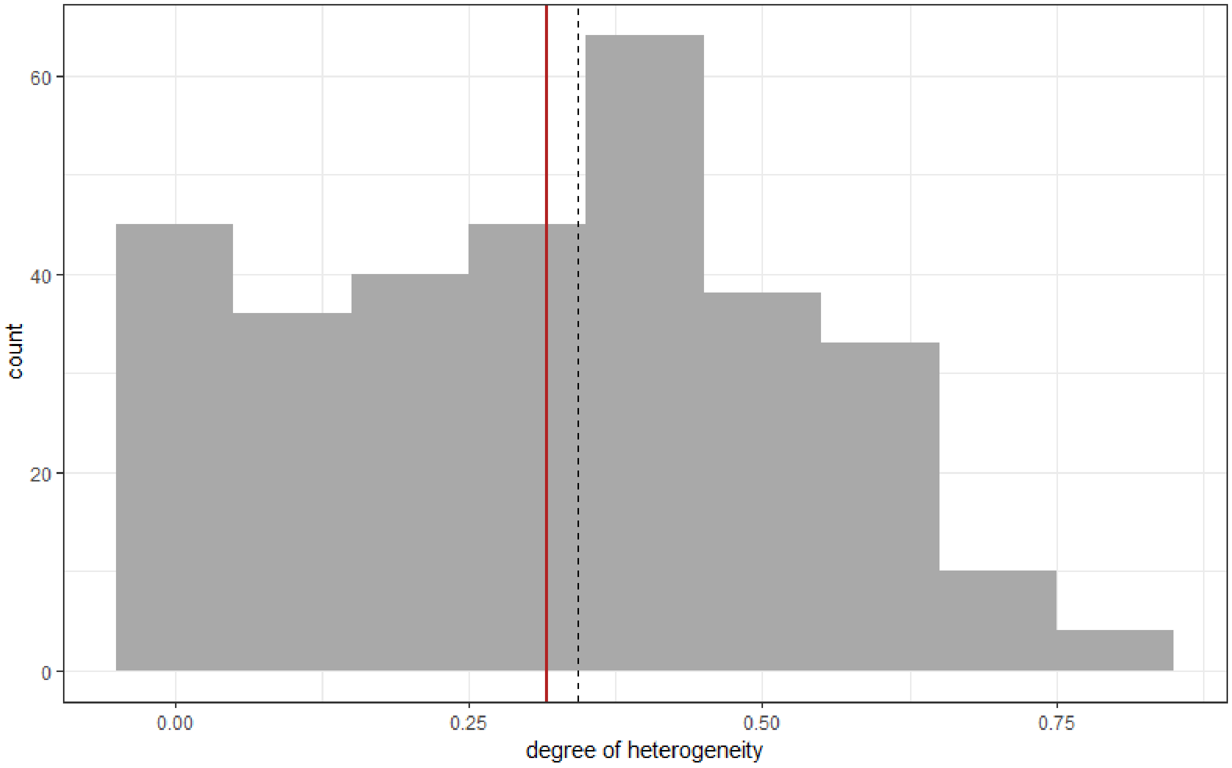

**Figure S2 | Link between public goods production and growth.** Depicted are the relationships between growth and the production of (a) biosurfactants, (b) biofilm components, (c) proteases, (d) pyoverdines, and (e) toxic compounds. All values are scaled relative to the corresponding values of our reference strain *P. aeruginosa* PAO1. Red lines indicate fits derived from robust linear models. Shaded areas (grey) are confidence intervals.

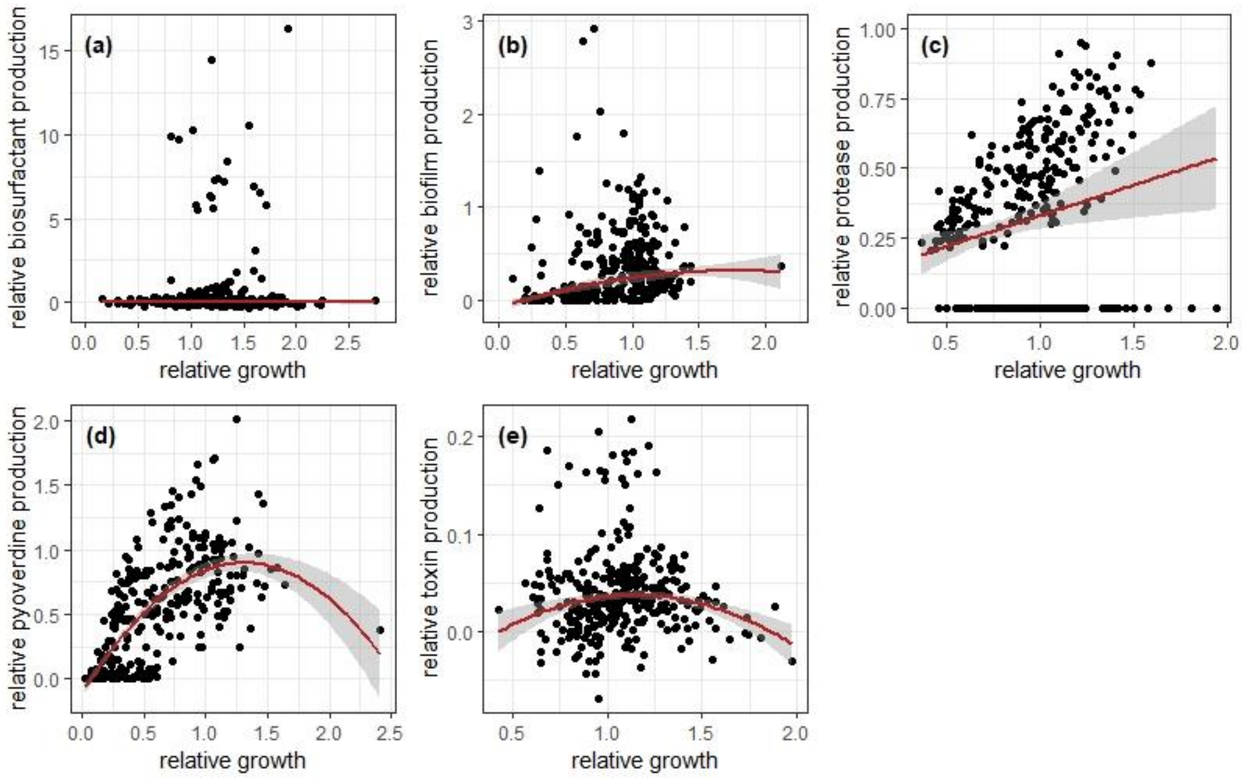
